# Disturbance regime changes leave long-lasting legacies on a microbial community’s composition and function

**DOI:** 10.64898/2026.06.09.731157

**Authors:** H. Inamine, L. Lear, A. Miller, S. Roxburgh, A. Buckling, K. Shea

**Affiliations:** Pennsylvania State University; University of Exeter; Commonwealth Scientific and Industrial Research Organization; Pennsylvania State University Main Campus: The Pennsylvania State University - University Park Campus

## Abstract

Mortality-inducing disturbances are important, ubiquitous drivers of community composition and function. Importantly, human activities and climate change are increasingly altering disturbance regimes. Most disturbance studies focus on the effects of current disturbance regimes, rarely considering those of historical regimes. However, recent theoretical work predicts that historical regimes can leave persistent legacies, modulating the community*’*s response to novel disturbances and invasive species. Here, we complement this theoretical approach using a model bacterial system that experienced disturbance regimes for ∼120 generations, followed by novel regimes and invasions for another ∼120 generations. Our results show persistent effects of historical legacies on disturbance-diversity relationships. Furthermore, some combinations of past and novel regimes promote invasion with increasing resident diversity, while others prevent it; legacies may explain conflicting diversity-invasibility relationships. These findings demonstrate the importance of historical legacies in disturbance-prone ecosystems, and underscore the challenges in predicting future community responses to disturbance regime changes.

## Introduction

Events that destroy biomass and release resources, i.e., disturbances, are ubiquitous and important drivers of community diversity (Connell 1978; Huston 1979; Petraitis *et al*. 1989; Pickett & White 1985; Sousa 1984). Disturbances may also affect the functions of a community, such as its robustness against invasions (Lockwood *et al*. 2008). Ecologists have empirically shown how disturbances affect diversity of plant (Biswas & Mallik 2010; Crawley 2004), animal (Bicknell & Peres 2010; Elton 1958), and microbial (Buckling *et al*. 2000; Lear *et al*. 2023) communities, and analyzed theoretical models to elucidate mechanisms that may be shared across diverse systems (Chesson & Huntly 1997; Levin & Paine 1974). Understanding how disturbances affect community diversity and functions is especially pressing as many communities face novel disturbances due to climate change (Newman 2019). Consequently, recent theoretical work has sought to understand how long-lasting effects of historical disturbances, or disturbance legacies, modulate how communities respond to future disturbances (e.g., (Miller *et al*. 2021)). Unfortunately, empirical work to examine the potential for these effects remains lacking. Here, we present empirical work to examine the potential for these effects.

Disturbances can take various forms such as fire, flood, antibiotics applications, and disease outbreaks. A key conceptual advance that synthesizes the effects of various disturbances is in characterizing them as regimes -- a set of spatiotemporal aspects of events such as frequency, intensity, duration, extent, timing, and pace (Walker 2011). Under this framework, different forms of disturbance (e.g., fire vs. flood) can be considered simultaneously, allowing for general insights to be drawn. However, communities under different regimes, for example different disturbance frequencies (i.e., how often disturbance events recur), may experience diverging diversity outcomes due to nonlinear effects of the mortality events on the dynamics of the surviving organisms (Miller *et al*. 2011). These patterns of diversity outcomes across communities under different disturbance regimes are called disturbance-diversity relationships. Much work has sought to characterize the general patterns of disturbance-diversity relationships across various aspects as well as systems (Mackey & Currie 2001). An underlying assumption in much of this body of work, however, is that disturbance regimes stay the same over time, or that the legacies of the historical regimes have been overwritten by ongoing disturbance regimes.

Recent work, however, has shown that the effects of past regimes may persist over many generations, long after communities experience novel regimes. In theoretical models, Miller and colleagues have demonstrated that these biotic legacies may persist for hundreds of generations even in the absence of evolutionary changes in the populations (Miller *et al*. 2021). The effects of regime changes are predicted to lead to complicated effects on the disturbance-diversity relationship: the diversity outcome of a community is a product of its past regime, the novel regime, and the interaction between the two. Similarly, there is a growing number of natural observations on how the dynamics of ecosystems may depend on past events, i.e., ecological memory may persist (Johnstone *et al*. 2016; Khalighi *et al*. 2022). However, experimental tests of these predictions are lacking because it is difficult to track the long-term dynamics of a community under regime change while exploring the possible combinations of the past and novel regimes. Here, we leverage the short generation time of the bacteria *Pseudomonas fluorescens* to analyze the persistent effects of disturbance legacies.

Disturbance regime changes may not only affect the diversity of a community, but also its function, such as its invasion resistance. A key hypothesis that explains the invasibility of a community is the diversity-invasibility relationship (Elton 1958), which posits that more diverse communities may better resist invasions because: 1) they deplete a variety of niches and therefore may have fewer available niches for the invaders (Herbold & Moyle 1986; MacArthur 1970); and 2) they are more likely to contain species that restrict successful colonization by the invaders, for example, competitors and consumers of the invaders (Elton 1958). While some empirical studies support the diversity-invasibility relationship (Case 1990; Knops *et al*. 1999; Levine 2000), others have shown the opposite pattern, where the less diverse communities better resist invasions (Davies *et al*. 2005; Stohlgren *et al*. 2003, 2006), perhaps because a diverse community may indicate a more resource-abundant ecosystem (Davis *et al*. 2000) or because diversity begets diversity (Madi *et al*. 2020; Palmer & Maurer 1997; San Roman & Wagner 2021). We hypothesize that differences in disturbance regimes may help explain these conflicting diversity-invasibility patterns, because disturbances can indirectly change the available niches for the invasive species by altering the resident diversity, while also directly reducing the propagules of the invaders (preventing invasion) as well as reducing the densities of the resident species (promoting invasions; (Hobbs & Huenneke 1992)). While the effects of disturbances on diversity and of diversity on invasions have been studied separately, less is known about how disturbances both directly and indirectly alter invasion outcomes. Furthermore, changes in disturbance regimes may additionally affect invasion success because the resident communities formed under the past regimes may be maladapted to the novel regimes; the transient dynamics that the communities experience under novel regimes may provide an opportunity for invaders to establish and grow. Communities experiencing disturbance regime changes may therefore be more prone to invaders. Changes in disturbance regimes can therefore complicate prediction of diversity-invasibility relationships because the causal relationships between disturbance regimes, community diversity, and invasion success are highly entangled.

Here, we address these knowledge gaps by using the bacterium *P. fluorescens* – an established model system for testing disturbance and invasion theories – to test how diversity is affected by historical disturbance regimes and by invasions. *P. fluorescens* rapidly diversifies into morphotype strains. Past work has shown that these morphotypes are differentially affected by the frequency of disturbances, which in turn affects the diversity and invasibility of this system (Buckling *et al*. 2000; Lear *et al*. 2022). Here, we adapted *P. fluorescens* to one of three disturbance frequency regimes for 8 days, followed by another regime with or without a mutant invader for an additional 8 days. We then quantified diversity and invasion success of the mutant, to test the hypothesis that historical disturbance regimes can leave a long-lasting legacy in the community. We found that, even after ∼120 generations under a novel regime, there is still an effect of the past regime on the diversity and the invasibility of the resident community. Furthermore, we show that the nature of the diversity-invasibility relationship depends on both past and novel regimes.

## Material and methods

### Bacterial cultures

Ancestral *Pseudomonas fluorescens* SBW25 was grown for 24 hours in King*’*s Medium B (KB). In a static condition, *P. fluorescens* rapidly diversifies into different morphotype strains that occupy different parts of liquid culture: the air-liquid interface (wrinkly spreader morphotype), the liquid column (smooth morphotype), or the bottom (fuzzy spreader morphotype) (Rainey & Travisano 1998). Sixty microliter of the culture was aliquoted to 18 sterile 20 mL microcosms containing 6 mL of KB, and the samples were cultured following the disturbance regimes detailed below. Following previous studies (Lear *et al*. 2020, 2022), a mutant *Pseudomonas fluorescens* SBW25-*lacZ* strain was used as an invader; the neutral *lac-Z* marker makes the mutant colonies visibly distinguishable from the resident wildtypes, due to blue pigmentation, on agar plates supplemented with X-gal (5-bromo-4-chloro-3-indolyl-β-D-galactopyranoside (Zhang & Rainey 2007)). SBW25-*lacZ* was cultured daily for 24 hours and the density was adjusted prior to invasion to an optical density at 600 nm of 0.1 with M9 salt solution (3 g KH_2_PO_4_, 6 g Na_2_HPO_4_, and 5 g NaCl per L). All samples were cultured in a static gravity convection incubator at 28 °C for the duration of the experiment.

### Disturbance regimes and invasions

Following previous studies (Buckling *et al*. 2000; Hall *et al*. 2012; Lear *et al*. 2022), we used *cell passage* as the disturbance in this system: a sample was vortexed for 30 seconds and 60 μL of it was transferred to a sterile microcosm containing 6 mL of KB (resulting in a random survivorship of 1%, or a disturbance intensity of 99%). Each sample was first passaged according to their disturbance frequency schedule for 8 days: passage every 1, 4, or 8 days (*“*past disturbance regimes*”*; *n* = 6 replicates per past disturbance regime; Figure *1*). On Day 8, all 18 communities were transferred into six sterile microcosms (108 total samples) that each followed 8 additional days of novel disturbance frequency schedule: passage every 1, 4, or 8 days, with or without daily invasions (*“*novel disturbance regimes*”*; *n* = 6 replicates of 9 combinations of past and novel disturbance regimes). Hereafter, we use → to signify how the disturbance regime changed (e.g., 1 → 8 signifies the change in disturbance regime from every 1 to every 8 days). Invaded samples were inoculated daily with 60 μL of the density-adjusted SBW25-*lacZ* culture (average invader CFU introduced = 7.2 ⨯ 10^5^ CFU) whereas the uninvaded samples were inoculated daily with 60 μL of M9 salt solution. Samples were archived in 25% glycerol mixture at -80 °C when they were passaged.

**Figure 1.**
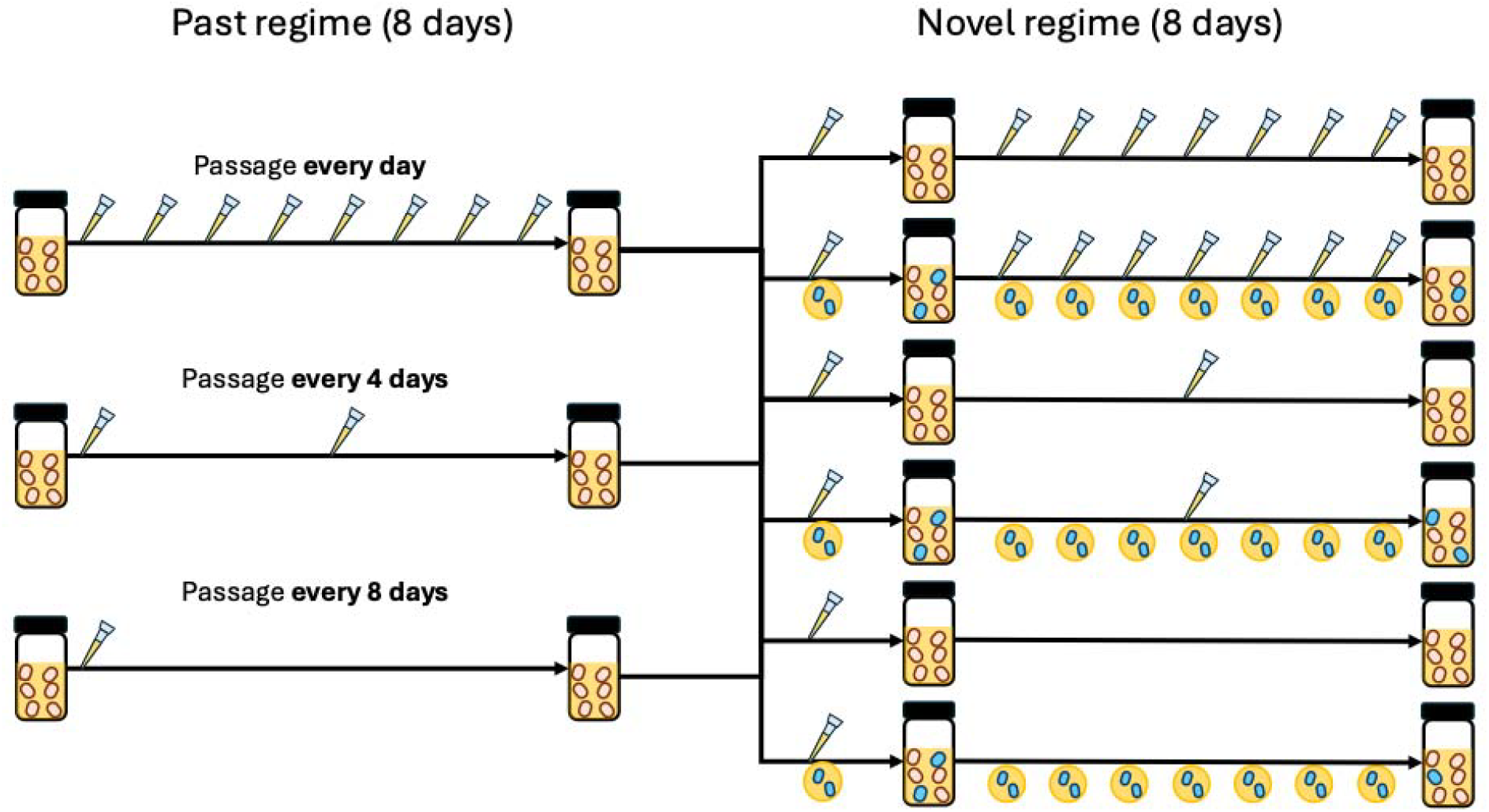
Conceptual diagram of the experiment. After 8 days of passage regimes (represented by different numbers of pipette tips), each microcosm was aliquoted into 6 fresh microcosms and underwent novel regimes with or without daily invasions (represented by blue bacteria cells). Samples were collected at each passage and on the final day of the experiment (Day 16).

### Statistical analyses

Resident and invader CFU densities of the SM, WS, and FS morphotypes and SBW25-*lacZ* were enumerated visually on KB agar plates supplemented with X-gal. The morphotype diversity of the wildtype strain (*“*resident diversity*”*) was calculated as the first-order effective number of morphotypes defined as α = *exp*(- ∑ *p*_*i*_ *log p*_*i*_), where *p*_*i*_ is the proportion of the *i*^th^ morphotype (Jost 2006). Community dissimilarity between the time points before (Day 8) and after (Day 16) novel regimes was calculated as *β*-1 = (*γ*/*a*) - 1, where *β* = exp(- ∑ *q*_i_ *log q*_i_) and *q*_*i*_ is the average proportion of the *i*^th^ morphotype across the two samples (Jost 2007).

A body of work on this system has established that the morphotype diversity shows a unimodal (i.e., quadratic) relationship with increasing passage frequency (Buckling *et al*. 2000; Hall *et al*. 2012). To assess the immediate effects of the passage regimes, we therefore first performed linear regression on the resident diversity of the samples collected on Day 8 after the past regimes, but prior to experiencing the novel regimes, using the model:

Equation 1

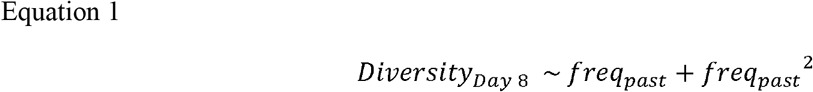

where *freq*_*past*_ is the frequency of the past regime (1, ¼, or 1/8). To assess the legacy effects of the past regimes, we performed linear regression on the resident diversity of the samples collected on Day 16 after the novel regimes, using the model:

Equation 2

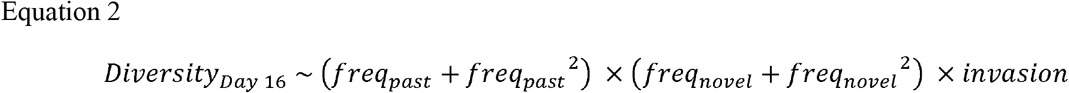

Equation 2 contains the direct effects of the past regimes, the novel regimes (*freq*_*novel*_), the invasions (*invasion*), and the interactions of these terms. Because we hypothesized that the morphotype diversity may exhibit a unimodal relationship with either the past or the novel regimes, we included quadratic terms for both regimes. We performed backward model selection (Johnson & Omland 2004) on this model, where we successively compare and refine the model to its sub-models based on AIC scores (Akaike 1974), to assess how the legacy effects interact with the novel regimes.

In addition to the communities*’* diversity profiles at different time points, we were also interested in understanding how the communities change and differ over time. We performed linear regression on community dissimilarity, using the model:

Equation 3

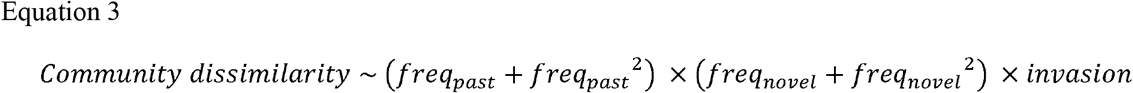

We performed backward model selection on this model to assess how disturbance regimes affect the magnitude of change.

To understand how the past and novel regimes affect invasion success, we fitted Poisson regression on the CFUs of the SBW25-lacZ invaders against the regimes using the model:

Equation 4

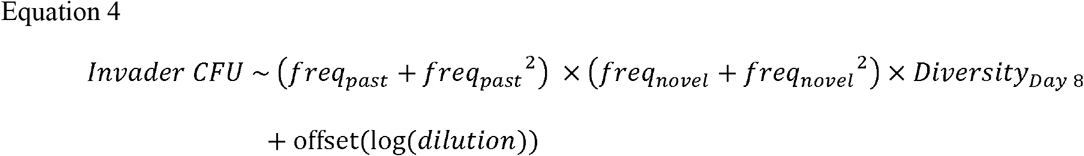

where the offset accounts for the dilution factors used to enumerate CFUs. First, to assess whether diversity affects invasion outcome, we constructed sub-models of Equation *4* that omit one or both of the quadratic terms. This resulted in differentially complex sub-models, and we compared each of their AIC scores with and without the *Diversity*_*Day 8*_ term. Second, we performed backward model selection on Equation *4* to assess how the legacies interact with the novel regimes.

Lastly, we performed a counterfactual analysis to assess whether ignoring regime changes could lead to misleading analyses or lower statistical power to detect ecological signals. To do so, we reanalyzed our data, focusing on treatments that experienced different past and novel regimes, yet assuming that the regimes were constant over the experimental timeframe. With this assumption, we recalculated the passage frequencies over 16 days (or equivalently, the total number of passages) instead of separately calculating them over the 8-day periods before and after the regime changes. For example, samples in the 8 → 4 treatment experienced 3 passages total (1 before and 2 after the regime change). We then performed ANOVA on the model

Equation 5

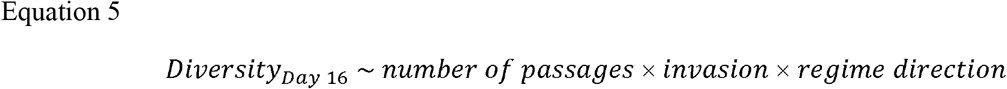

where *regime direction* indicated whether the disturbance regime increased or decreased over the experiment. All statistical analyses were performed in R version 4.3.1 (R Core Team 2022).

## Results

Consistent with prior work (Buckling *et al*. 2000; Hall *et al*. 2012), passage frequency regimes of every 1, 4, or 8 days resulted in communities with a unimodal diversity pattern with increasing frequencies (Figure 2A; Adjusted R^2^ = 0.53, *F*(2, 15) = 10.54, *p* = 0.0014). Specifically, diversity increased with the past frequency (*freq*_*past*_ effect size = 2.76, *p* = 0.03) but decreased with its quadratic term (*freq*_*past*_^2^ effect size = -2.69, *p* = 0.02), such that intermediate frequencies saw the most diverse populations, followed by low frequencies, then high frequencies, before the communities were subjected to novel regimes. Given these effects on the diversity outcomes, we next tested whether the past regimes leave lasting legacies on the communities that could be detected after 8 days, or ∼120 generations, after the regime change. Backward model selection on Equation 2 showed that the best model to explain the resident diversity 8 days after the regime changes is

**Figure 2.**
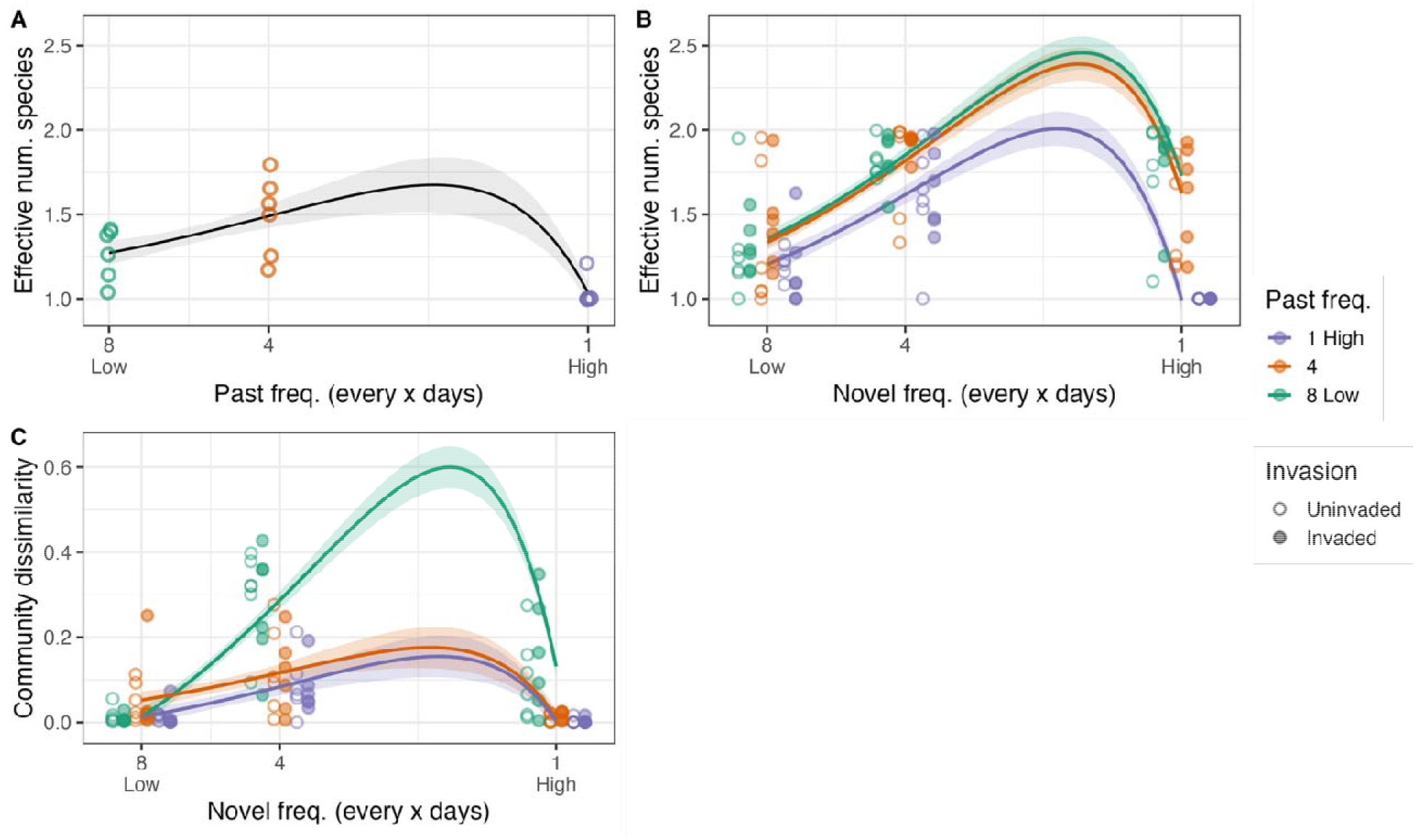
Effects of passage frequencies on the morphotype diversity of *Pseudomonas fluorescens* strains (A) before and (B) after the disturbance regime change, and (C) the dissimilarity between the two time points. The curves and their envelopes are the predicted estimates and the standard errors from the best fit models, and the individual points are the data (colors: past regimes; unfilled: uninvaded, filled: invaded).

Equation 6

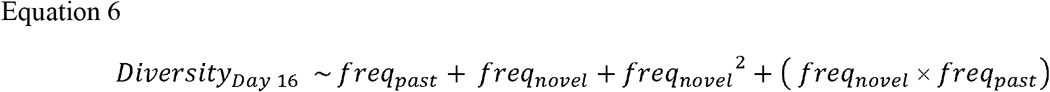

The communities displayed a unimodal diversity pattern similar to that on Day 8, but this pattern on Day 16 was driven mostly by the novel frequencies (Figure *2*B; Adjusted R^2^ = 0.57, *F*(4, 103) = 36.01, *p* < 2.2 ⨯ 10^−16^): diversity increased with the novel regime (*freq*_*novel*_ effect size = 5.85, *p* < 10^−13^) but decreased with its quadratic term (*freq*_*novel*_^2^ effect size = -4.74, *p* < 10^−12^). Legacies from the past regimes did not have a statistically significant direct effect (*freq*_*past*_; *p* = 0.41), but they interacted with the novel regimes (*freq*_*novel*_ : *freq*_*past*_ effect size = -0.76, *p* < 10^−5^). Furthermore, Equation 6 implies that, for a single value of *freq*_*novel*_, diversity decreases monotonically with increasing past regime. This can be seen in Figure *2*B, where the peak of the disturbance-diversity relationship decreases with increasing frequency. While the magnitude of the interactive term was not large enough to alter the unimodal diversity pattern, the legacies from the past regimes nonetheless persistently and quantitatively affected disturbance-diversity relationship under the novel regimes.

Surprisingly, invasion had no statistically significant effect on the resident diversity in our experiment, even though they compete heavily for resources with the wildtype strain: all terms associated with invasion were absent in the final model in Equation 6. To assess whether invasion affected resident diversity under certain combinations of past and novel regimes, we additionally performed ANOVA on resident diversity but treated past and novel frequencies as unordered factors instead of continuous variables. This analysis showed qualitative agreement with the original results: the past and novel regimes affected diversity (*p <* 10^−3^ for their main and interactive effects), but invasion did not (*p* > 0.3 for all terms involving invasion).

While the above analyses demonstrated the effects of disturbances on diversity at the end of the novel regimes, they did not assess how much the communities changed over the course of the regimes. We performed backward model selection on Equation *3* in order to do so, which resulted in

Equation 7

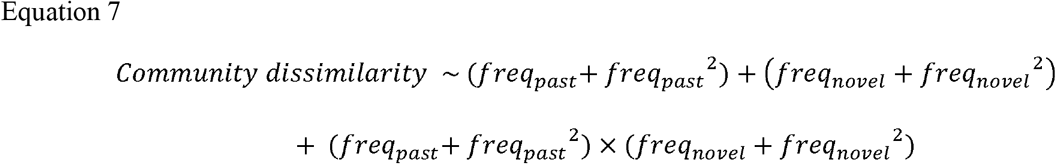

where community dissimilarity measures the dissimilarity between a community before (Day 8) and after (Day 16) the novel regime (0: identical; 1: completely different). All terms with invasion were dropped from the model, indicating that the interactions with the invaders did not affect community dynamics (Adjusted R^2^ = 0.59, *F*(8, 99) = 20.15, *p* < 2.2 ⨯ 10^−16^). Community dissimilarity showed unimodal patterns against both the past (effect size for *freq*_*past*_ = 3.61, *freq*_*past*_^2^ = -2.93, *p* < 10^−3^ for both) and novel (effect size for *freq*_*novel*_ = 6.36, *freq*_*novel*_^2^ = -5.32, *p* < 10^−9^ for both) regimes. Furthermore, the two regimes interacted (effect size for *freq*_*past*_ : *freq*_*novel*_ = -27.85, *freq past*^2^ : *freq*_*novel*_ = 22.34, *freq*_*past*_ : *freq*_*novel*_^2^ = 22.87, *freq*_*past*_^2^ : *freq*_*novel*_^2^ = -18.32; *p* < 10^−4^ for all), resulting in a complex pattern of changes in the community over time. For example, communities that experienced disturbances every 8 days in the past regimes experienced large changes under novel regimes of every 4 and 1 day, but not under the novel regime of every 8 days (Figure *2*C).

Given the lasting effects of the past regimes on the dynamics and the outcomes of resident diversity, we hypothesized that the disturbance legacies may similarly alter invasion outcomes. First, we found that, while both past and novel regimes are necessary to explain invasion outcomes, they are not sufficient; across differentially complex sub-models of Equation *4*, incorporating diversity always improved model performances (*Diversity*_*Day9*_ lowered AIC scores for every model by > 2). Second, backward model selection on Equation 4 showed that the best statistical model to explain invasion outcome is

Equation 8

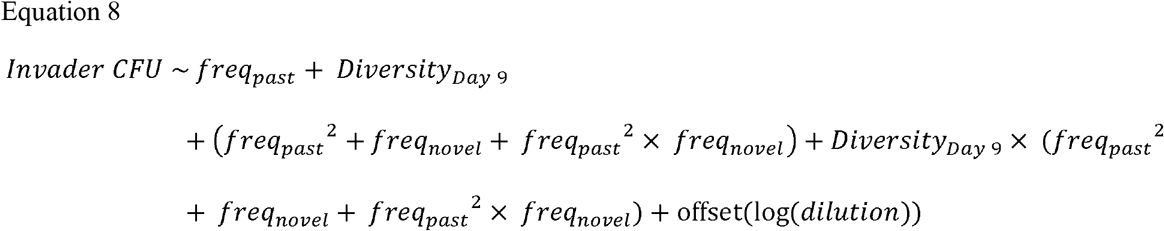

Invader density increased with increasing diversity (*Diversity*_*Day9*_ estimate: 4.4, *p <* 10^−3^), but it was also directly affected by the past and novel disturbance frequencies (estimates: *freq*_*past*_ *=* -9.1, *freq*_*past*_^*2*^ *=* 20.0, *freq*_*novel*_ *=* 13.4, and *freq*_*past*_^*2*^ : *freq*_*novel*_ = -27.0; *p* < 0.05 for all terms). Furthermore, both past and novel disturbance regimes indirectly determined invasion outcomes, as they interacted with and altered the effects of diversity on invasions (estimates: *Diversity*_*Day9*_ : *freq*_*past*_^*2*^ *=* -11.2, *Diversity*_*Day9*_ ⨯ *freq*_*novel*_ *=* -9.4, and *Diversity*_*Day9*_ : *freq*_*past*_^*2*^ : *freq*_*novel*_ = 23.8; *p* < 0.05 for all terms). Taken together, the invader density decreased with diversity when the communities experienced frequent disturbances in either the past or the novel regimes, but not both. Under other conditions, it increased with diversity (Figure *3*). Third, because the model in Equation 8 is fairly complex, we performed an additional analysis to assess the sensitivity of our results to this particular model structure. We picked the model that had the lowest AIC score from our prior analysis of the differentially complex sub-models, and inspected how the diversity-invasibility relationship changes under different combinations of the regimes. The resulting model was also the simplest one: *Invader CFU ∼ freq*_*past*_ ⨯ *freq*_*novel*_ ⨯ *Diversity*_*Day9*_. This analysis showed qualitative agreement with the original results: the invader density increased with diversity under the same regime combinations, and decreased under others.

**Figure 3.**
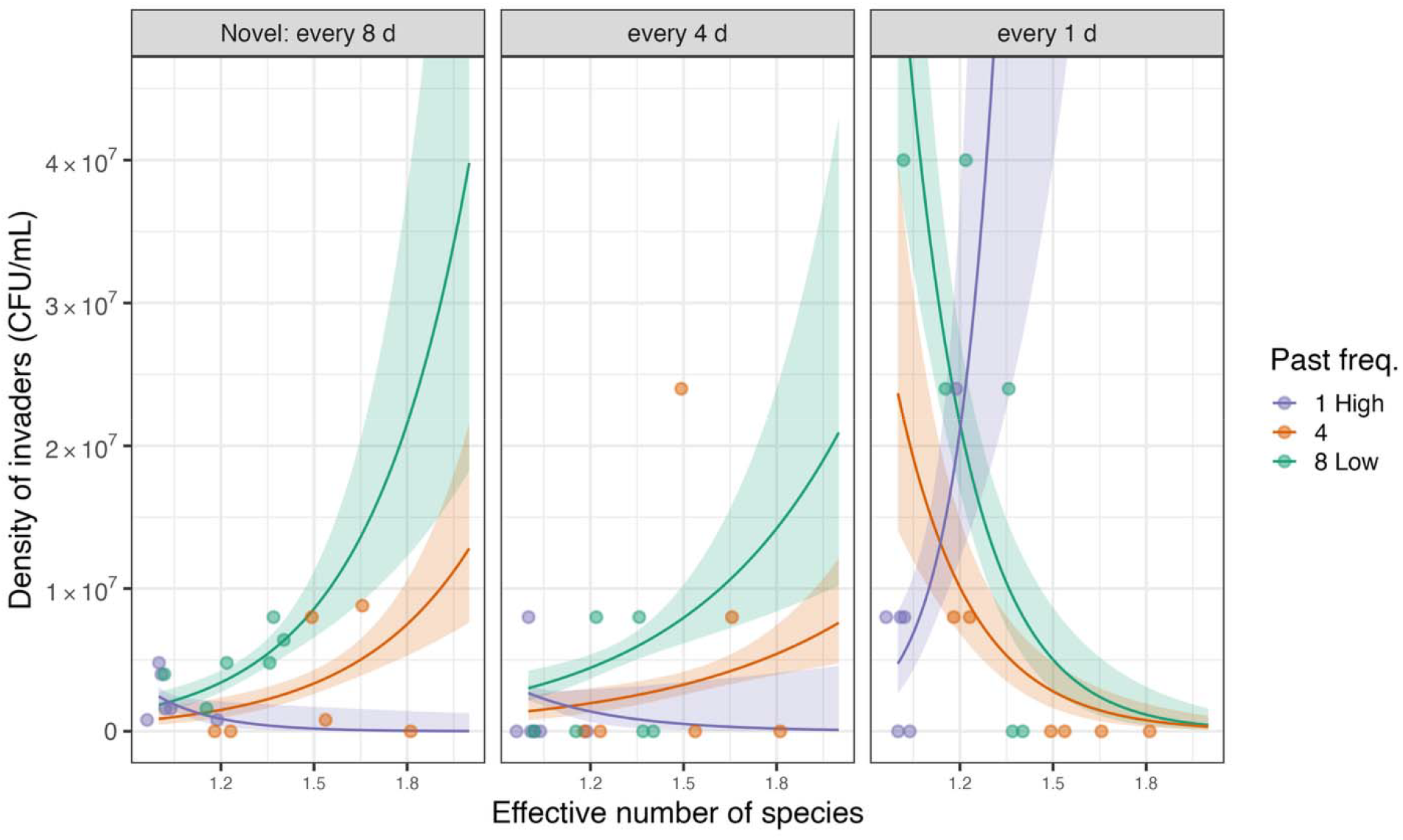
Effects of passage regimes on diversity-invasibility relationships. Each panel shows the effects of diversity on invader density under each novel regime (colors: past regimes). The curves and their envelopes are the predicted estimates and the standard errors from the best fit model, and the individual points are the data.

While the analyses above show how the past regimes, novel regimes, and the resident community structures jointly affect invasion outcomes, they do not assess whether the transient dynamics of the community structures under novel regimes may provide an opportunity for the invaders to grow. To test this, we analyzed the model focusing on whether regime change alone can help explain the differences in invasion outcomes. We found that, while there were significant differences in the invasion outcomes under specific regime changes (estimated marginal means for 8 → 1 vs. 8 → 8 is 4.8 and for 1 → 8 vs. 1 → 1 is 0.28; *p* < 0.05 for both contrasts), there were no overall patterns of invasibility (*p* = 0.25 for a Poisson regression that analyzed treatments with different vs. constant regimes). Instead, we found that the community dissimilarity index, which measures resident community turnover, can help explain invasion outcomes (*Community dissimilarity* estimate: 3.53, *p* < 0.01). Communities that underwent drastic changes under regime changes are therefore more susceptible to invasion.

Our study demonstrates the importance of a regime change as an event in its own right that must be explicitly considered: a change in disturbance regime is itself a disturbance (Miller *et al*. 2018). Ignoring regime change – the fact that the frequency increased or decreased -- may therefore lead to faulty or weaker inferences. In order to test this hypothesis, we re-analyzed our diversity data by the total number of disturbances over the whole experimental timeframe (16 days) as well as the direction of disturbance frequency change. If regime change is important, then the direction of the regime change should have a significant effect. ANOVA on Equation *5* showed that the direction is indeed significant (*regime direction F*(1, 60) = 24.4, *p* < 10^−5^) and also interacted with the total number of disturbances (*number of passages* : *regime direction F*(2, 60) = 8.58, *p* < 10^−3^). Furthermore, the model in Equation *5* performed better than the model without *regime direction* terms (ΔAIC = 26). Thus, ignoring disturbance regime change led to weaker inferences and misses identifying an important factor influencing community diversity.

## Discussion

Disturbances are an important determinant of community structure and function, and ecological systems have long adapted to them through both ecological and evolutionary processes. Changes to the disturbance regimes with which communities are familiar may therefore perturb the communities away from their norms, leading to loss of species, loss of ecosystem services, and establishment of invasive species. Not surprisingly then, there is a growing interest in understanding how regime changes might affect the structure and the function of ecological communities; disturbance regimes are changing globally. However, it has been difficult to test these questions in natural systems because of the time needed for legacies to build and persist. Here, we leverage the short generation times of *P. fluorescens*, a microbial system that has been particularly fruitful for testing disturbance theories, to understand how historical disturbances can leave long-lasting legacies. Our results support key hypotheses generated by the theoretical models of Miller and colleagues (Miller *et al*. 2021). First, the models predicted that the legacy effects of historical disturbances can endure for a surprisingly long time. In our experiments, the bacteria underwent novel regimes for ∼120 generations, yet they still persistently exhibited the effects of the past regimes on both the diversity and the invader density. While community diversity had a unimodal pattern against the novel frequencies, the communities that experienced higher past disturbance consistently had lower diversity (Figure *2*B). Furthermore, our empirical work demonstrates that legacy effects can be incredibly important, and failing to consider them can lead to misinterpretation.

Second, the theoretical models predicted that the changes in invasion outcomes under a regime change may not mirror the changes in diversity of the resident species. Some past empirical studies have shown support for diversity-invasibility hypothesis, where an increase in resident diversity is posited to lead to a decrease in invasibility, but others have shown the opposite pattern: an apparent paradox (Fridley *et al*. 2007; Levine & D*’*Antonio 1999; Smith & Côté 2019). In our experiment, we observed both patterns to occur depending on the combination of the past and novel regimes (Figure *3*), potentially allowing us to explain this paradox. Specifically, we observed negative relationship between invader density and diversity if one, but not both, of the regimes consisted of a daily (i.e., high frequency) disturbance. In all other regime combinations, we observed the positive relationship. The pattern of Figure *3* suggests that there may even be a situation where invader density does not depend on resident diversity, as the positive relationship changes to negative. While previous studies have proposed different mechanisms that link diversity to invasibility (Levine & D*’*Antonio 1999), our results suggest that past disturbance regime may interact with the novel regime to quantitatively and qualitatively alter diversity-invasibility relationships even in the same ecological system: legacies of historical processes may be the missing factor needed to provide a unified understanding of how diversity can both ward off and encourage invasion in different contexts. Furthermore, our results address the importance of analyzing the combined direct and indirect effects of disturbances on invasibility. While researchers have studied the effects of disturbances on diversity (Connell 1978; Huston 1979), diversity on invasibility (Levine & D*’*Antonio 1999), and disturbances on invasion (Lembrechts *et al*. 2016; Roxburgh *et al*. 2004) separately in empirical and theoretical systems, what we now clearly need is a framework to understand how the effects of disturbances propagate through these tangled pathways. Our study provides a blueprint for such a framework.

There are a number of complementary ecological and evolutionary mechanisms that could help explain how regime changes affect invasion outcomes. First, the resident communities undergoing a shifting disturbance regime may be in flux, reducing the likelihood of them imposing dominance effects (disproportionately increasing invasion resistance by, for example, competitively dominating a resource; (Hodgson *et al*. 2002)) or priority effects (a fitness advantage from occupying a niche first (Fukami 2015)) that restrict colonization. The transient dynamics that communities experience as they relax back into novel steady states may therefore provide an opportunity for the invaders. Second, regime changes may leave evolutionary legacies in the community, as the organisms adapted to past regimes face different evolutionary forces under novel regimes (Bell 2017). Maladapted life-history traits may render the communities susceptible to invasions. Both ecological and evolutionary mechanisms are likely at work in our experimental system, as the short generation time provides ample opportunities for eco-evolutionary dynamics. While we did not measure the specific life-history traits of the bacteria, our results suggest that maladaptation alone cannot explain the invasion outcomes, as residing in a constant regime did not improve invasion resistance. Under a constant regime, we might expect that the morphotypes are pre-adapted to the past regime, and therefore more resistant to invaders in a regime that remains the same. Instead, we found that communities undergoing large structural fluxes are more prone to invasions. These two lines of evidence suggest that integrated approaches are necessary to gain predictive insight. Our work also suggests the need to expand current theoretical work on disturbance to include both ecological and evolutionary adaptation to disturbance regimes.

Past disturbance affected the community diversities at the end of the experiment, but it also altered how the communities change under novel regimes. Even though the communities that were disturbed every day had the least diversity (Figure *2*A), they were also the ones that changed the least under the novel regimes (Figure *2*C). The communities that were disturbed every 8 days in the past changed the most, even though they had intermediate amount of diversity. These results suggest how future diversity outcomes (in our experiment, Day 16) may not only be a function of the starting community diversity (Day 8), but also a product of historical regimes that could leave biotic and abiotic legacies in the communities. We also note that, while 1 → 1 and 8 → 8 showed very little changes, communities in 4 → 4 were different between Days 8 and 16. It*’*s possible that the 4-day regime may involve other mechanisms, but this hypothesis should be explored further in future experiments with a wider range of frequencies.

Lastly, our results highlight the importance of explicitly considering regime changes as disturbances in their own right. The direction of the regime change (i.e., increasing or decreasing frequency) left a clear difference in the resident diversity (Figure *4*). Our work shows that failing to consider the changes in disturbance frequency, and mischaracterizing the regime as a constant one, could lead to the wrong interpretation of how disturbances affect diversity. Regime changes may be a common occurrence if, for example, disturbances have temporal autocorrelation and are *“*clumped in time*”* (e.g. (Garrison *et al*. 2012)). Considering past regimes and regime changes is especially challenging because of their spatial and the temporal scopes: changes in disturbance regimes are happening globally and the legacy of the past regime could linger for hundreds of generations. Overcoming these challenges empirically may require longer and finer scale data to capture the past regime as well as to identify regime changes. Our rapid, tractable empirical approach provides important information for larger, slower and more complex ecological systems. Emerging theoretical work on transient and nonequilibrium dynamics (Hastings 2001; Hastings *et al*. 2018) may also provide some avenues to tackle these challenges.

**Figure 4.**
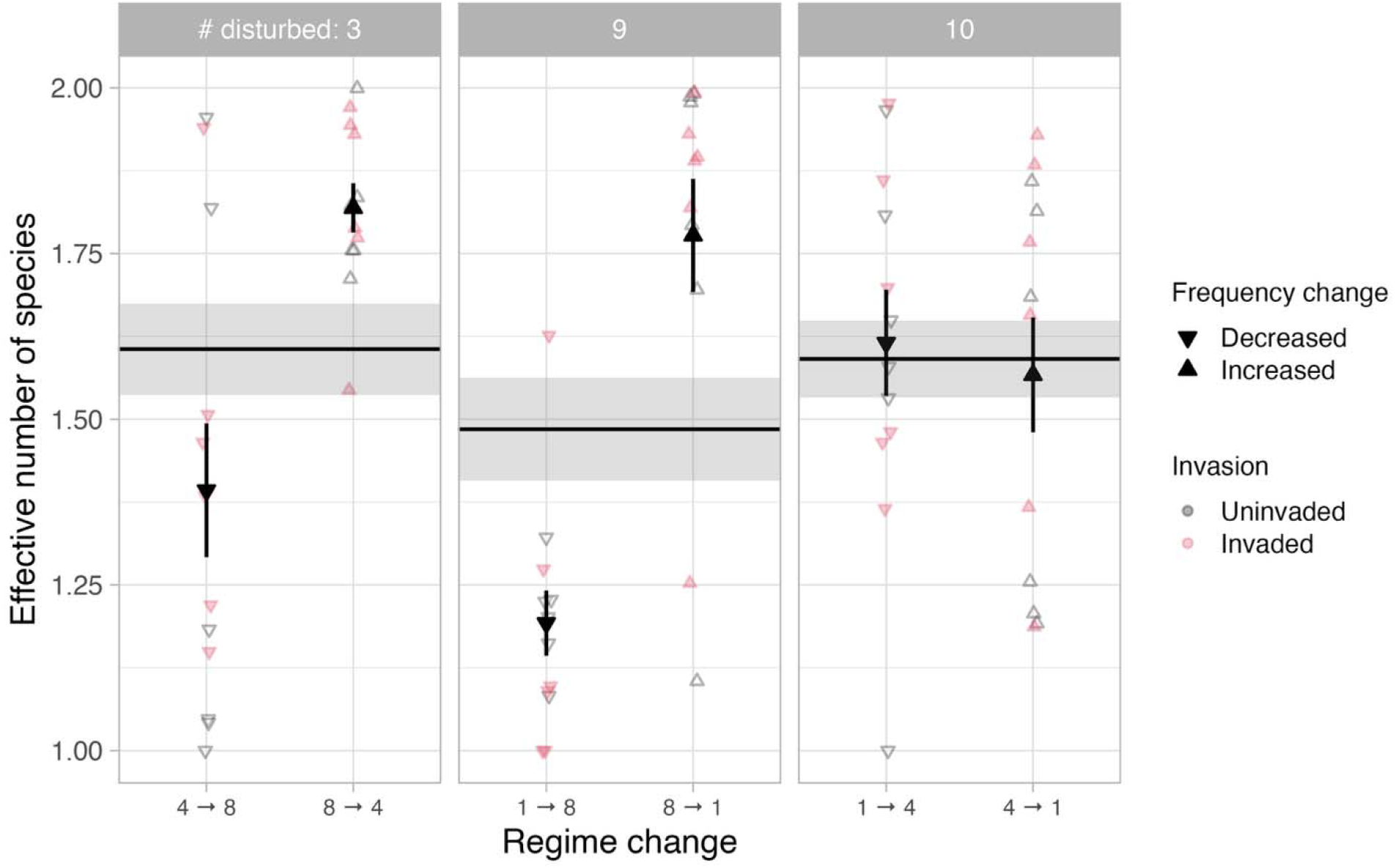
Counterfactual analysis on the effects of the total number of disturbances (3, 9, or 10 disturbances across both past and novel regimes) and invasion (filled: invaded; empty: uninvaded) on the diversity of the communities. We analyzed these data by considering the direction of the disturbance frequency change (down-pointing triangle: decreased frequency; up-pointing triangle: increased frequency). The horizontal lines and the envelopes are the means and the standard errors if the regime changes were ignored.

While the present work has focused specifically on how disturbance legacy affects invasion in an experimental bacterial system, our results suggest far-reaching consequences of legacy that may be applicable broadly to other ecological systems, processes, and applications. First, there is a pressing need to understand how disturbances promote or prevent invasions as we seek to manage microbial communities. Microbial biosecurity threats, such as pathogen spillover and environmental contamination, as well as microbiome engineering, such as microbiome restoration and agricultural bio-inoculation, are essentially about invasions in microbial communities (Jack *et al*. 2021; Ladau *et al*. 2025). Disturbances are ubiquitous in microbial communities, e.g., antibiotics that disrupt pathogens and microbiomes (Blaser 2016; Eckert *et al*. 2019), but these interventions have rarely been analyzed through the lens of disturbance and invasion theories (but see (Baker *et al*. 2018; Campbell *et al*. 2022)). Our work highlights the potential benefits of doing so, while underscoring the importance of disturbance legacy for prediction of invasion outcomes. Second, the legacy effects of various regime changes (e.g., prolonged drought, increasing hurricane intensity and frequency, longer fire seasons) could potentially affect conservation and management of invasive species in any ecological community. Furthermore, we hypothesize that the effects of legacy are not limited to diversity and invasion; they are likely multifaceted, potentially affecting other community functions such as productivity, nutrient cycling, and regulation of the abiotic environment. Our results highlight the urgent need to take historical context into account when analyzing ecological data, both for basic understanding of natural systems as well as for managing ecosystems of all types.

## Acknowledgements

This material is based upon work supported by the National Science Foundation under Award No. DMS-2347200 (HI and KS). LL thanks the UKRI FLF award MR/V022482/1.

